# The Anna Karenina model of β cell maturation in development and their dedifferentiation in type 1 and type 2 diabetes

**DOI:** 10.1101/2021.02.16.431507

**Authors:** Sutichot D. Nimkulrat, Zijian Ni, Jared Brown, Christina Kendziorski, Barak Blum

## Abstract

Loss of mature β cell function and identity, or β cell dedifferentiation, is seen in all types of diabetes mellitus. Two competing models explain β cell dedifferentiation in diabetes. In the first model, β cells dedifferentiate in the reverse order of their developmental ontogeny. This model predicts that dedifferentiated β cells resemble β cell progenitors. In the second model, β cell dedifferentiation depends on the type of diabetogenic stress. This model, which we call the “Anna Karenina” model, predicts that in each type of diabetes, β cells dedifferentiate in their own way, depending on how their mature identity is disrupted by any particular diabetogenic stress. We directly tested the two models using a β cell-specific lineage-tracing system coupled with RNA-sequencing in mice. We constructed a multidimensional map of β cell transcriptional trajectories during the normal course of β cell postnatal development and during their dedifferentiation in models of both type 1 diabetes (NOD) and type 2 diabetes (*BTBR-Lep^ob/ob^*). Using this unbiased approach, we show here that despite some similarities between immature and dedifferentiated β cells, β cells dedifferentiation in the two mouse models is not a reversal of developmental ontogeny and is different between different types of diabetes.

## Introduction

Insulin-secreting pancreatic β cells are essential for maintaining blood glucose homeostasis, and their loss or dysfunction underlies all types of diabetes mellitus. In type 1 diabetes (T1D), β cells are targeted by an autoimmune attack. In type 2 diabetes (T2D), β cells fail due to work overload and a toxic metabolic environment brought about by obesity and peripheral insulin resistance. In recent years, it has become clear that not all β cells are permanently lost in either type of diabetes. Instead, chronically stressed β cells lose their functionally mature phenotype and shift to a dysfunctional state in a process called dedifferentiation. Such β cell dedifferentiation is seen in humans (1–6) as well as in murine models of both T1D and T2D (7, 8). The progression to overt diabetes can be prevented if diabetic β cell stress is alleviated in time, before the functionally mature β cell mass is lost (9, 10). Thus, drugs that work by directly reversing or preventing β cell dedifferentiation are critically needed (11, 12).

The term “β cell dedifferentiation” to describe the loss of mature β cell phenotype was first coined over two decades ago (13, 14). However, what exactly constitutes “dedifferentiated β cells” remains debated (15). Previously, it was proposed that β cells in diabetes dedifferentiate in the reverse order of their normal developmental ontogeny (8). This model predicts that dedifferentiated β cells resemble β cell progenitors (Figure 1, top). An alternative model suggests that β cell dedifferentiation is a stress type-specific process caused by disruption of specific gene regulatory networks by the diabetogenic environment, thus resulting in a stress-type specific loss of functional maturity, without assuming a “true” β progenitor cell identity (16). This model, which we call the Anna Karenina model (based on the opening sentence in Tolstoy’s novel by the same name, “All happy families resemble one another, each unhappy family is unhappy in its own way” (17)), predicts that, in each type of diabetes, β cells will lose their mature phenotype in a unique manner, depending on how their genetic network is perturbed by a particular diabetogenic environment (Figure 1, bottom).

**Figure 1.**
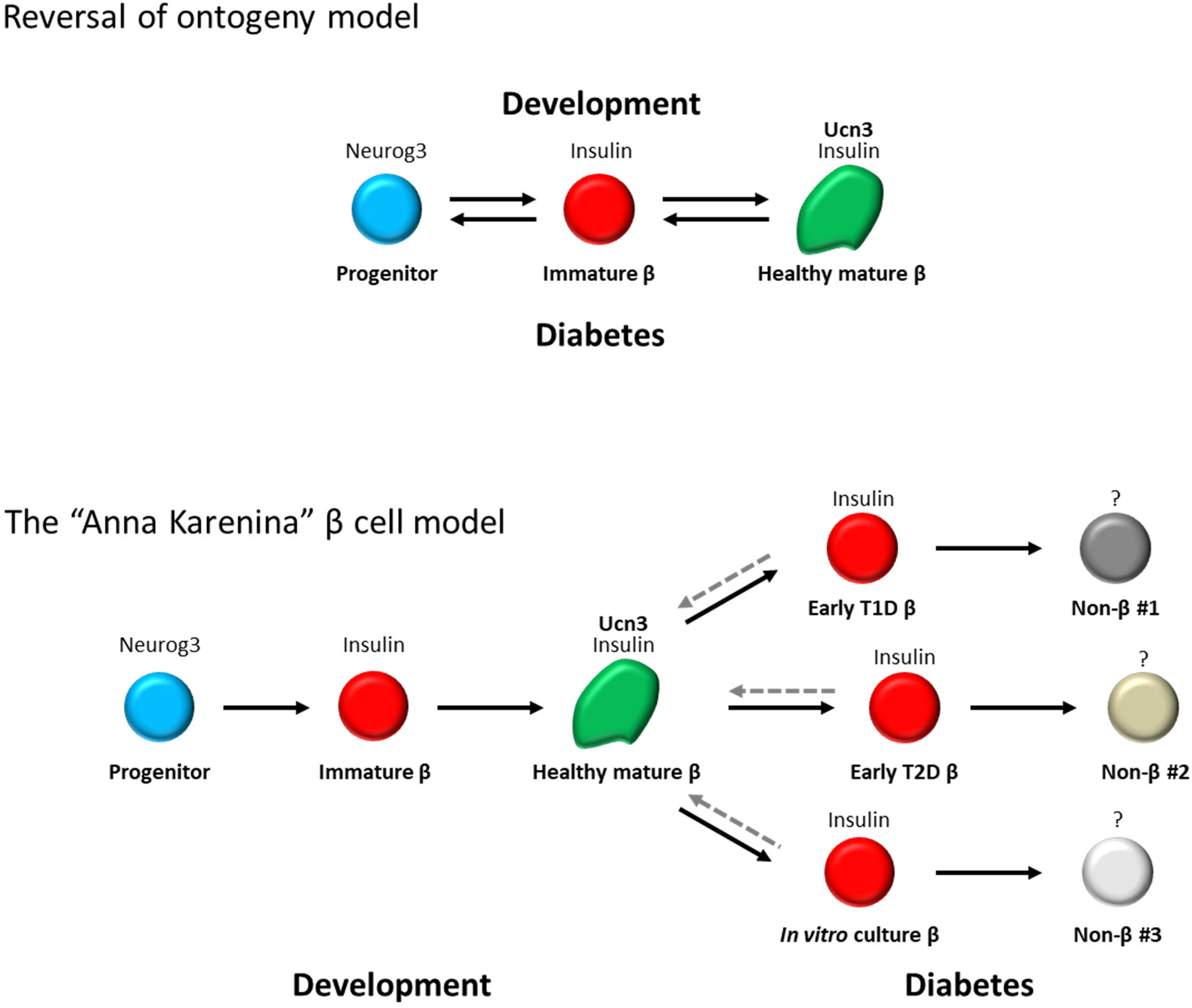
Two models for β cell dedifferentiation in diabetes. **Top:** The reversal of ontogeny model predicts that β cells dedifferentiate in diabetes in a reverse order of their normal ontogeny during development. **Bottom:** The “Anna Karenina” model predicts that in each type of diabetic stress, β cells lose their mature identity in a different way and take on different dedifferentiated identities.

Here, we test the Anna Karenina model of β cell dedifferentiation in diabetes. Specifically, we test whether under different types of diabetic stress, dedifferentiated β cells resemble one progenitor state, or if each type of diabetes produces β cells that are dedifferentiated in their own way. We do so by elucidating how the transcriptional landscape of β cells changes during their maturation in normal development, and their dedifferentiation in different types of diabetes, using a β cell-specific lineage-tracing system in mice. This approach enables us to follow β cells during both the normal course of their development and during their dedifferentiation in diabetes, and allows for direct, unbiased comparison between the gain of β cell maturation in development and the ways it is lost upon different types of diabetogenic insult.

## Results

### Transcriptional relationships between β cell maturation in postnatal development and their dedifferentiation in different types of diabetes

To test the transcriptional relationship between β cell maturation and their dedifferentiation in different types of diabetes, we used our previously reported murine β cell-specific lineage-tracing system (18, 19). This system is made by crossing mice transgenic for *Insulin2-Cre* with mice carrying a floxed reporter of histone H2B fused to mCherry (*Rosa26-lox-stop-lox-H2BmCherry*). In this system, any cell that had ever expressed the *Insulin* gene is permanently marked with nuclear mCherry. This reporter mouse line thus enables us to isolate and investigate β cells through development and functional maturation, as well as through the progression of diabetes, using a single-platform method. We crossed this system into the non-obese diabetic (NOD) model of autoimmune T1D and into the BTBR-*Lep^Ob/Ob^* (BTBR-*Ob/Ob*) model of obesity-related T2D. We FACS-purified lineage traced β cells from healthy mice during postnatal development, through adulthood, and during the progression to diabetes in the different models. We next subjected the samples to whole-genome RNA-sequencing. We thus generated gene expression data from four time points during β cell development and maturation (E18.5, P1, P7 and P10), as well as healthy adult mice and diabetic mice (defined by having fed blood glucose levels >300mg/dL).

We performed unsupervised bottom-up hierarchical clustering of the samples based on the top 15% most variable genes, using Spearman’s correlation as the distance metric (Figure 2). This method identified three large clusters (“development”, “healthy adult”, and “diabetic”). Importantly, wildtype (*WT)* samples (ICR genetic background) and non-diabetic *Ob/+* samples (BTBR genetic background) clustered together, without apparent separation between them, confirming that our method correctly distinguishes between the disease conditions, and not between genetic backgrounds. Interestingly, three of the NOD non-diabetic samples clustered together with the healthy adult samples, and four of the NOD non-diabetic samples clustered with the diabetes samples, suggesting that transcriptional changes related to β cell stress can be detected before the increase in blood glucose in these mice.

**Figure 2.**
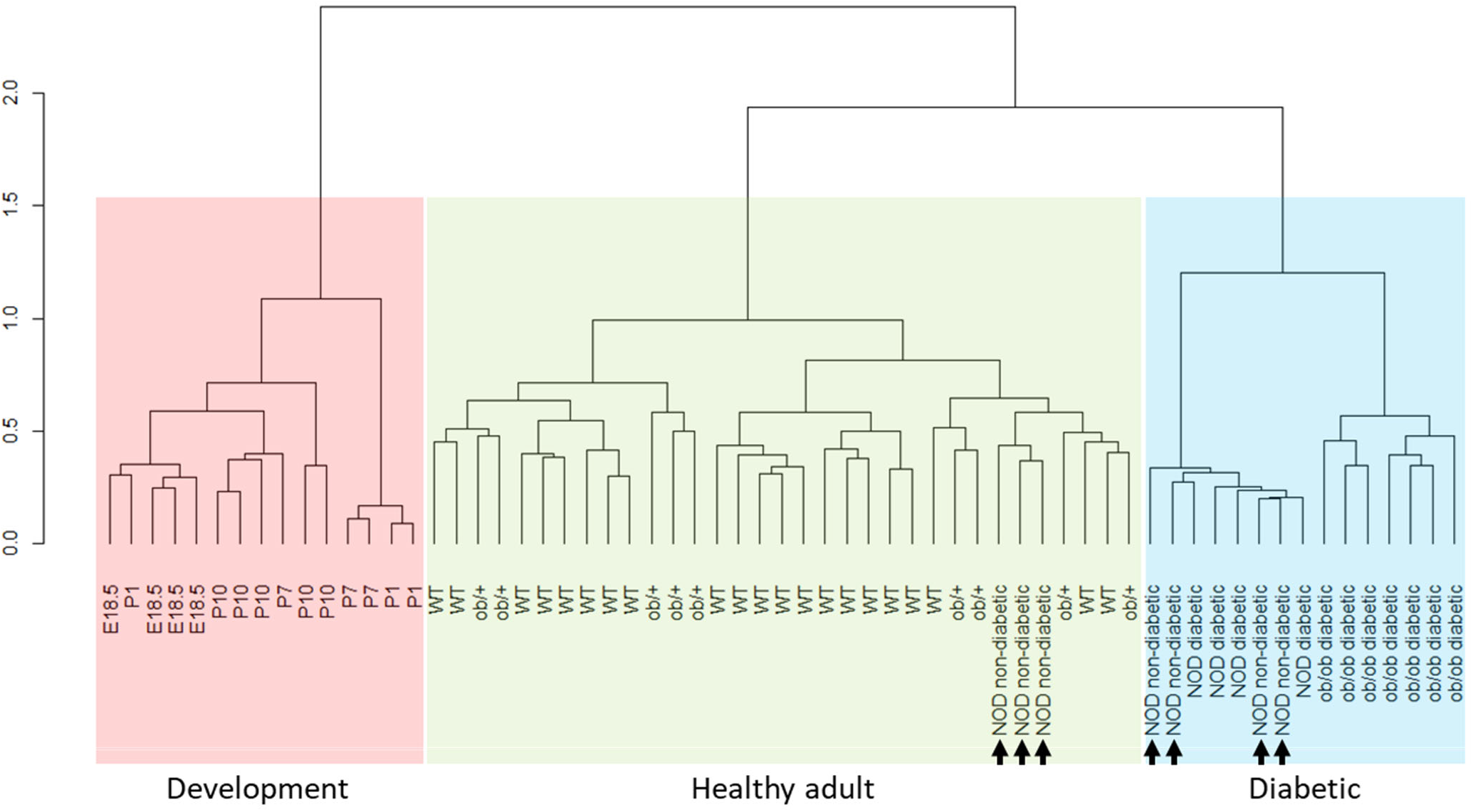
Unsupervised hierarchical clustering of β cell transcriptomes during development and dedifferentiation. Unsupervised bottom-up hierarchical clustering of the samples based on the top 15% most variable genes using Spearman’s correlation as the distance metric is shown. Three independent clusters are identified: development (red), healthy adult (green), and diabetic (blue). Adult *WT* and non-diabetic *Ob/+* samples cluster together, confirming that the method correctly distinguishes between the diabetes stages and not between genetic backgrounds. Three of the NOD non-diabetic samples cluster together with the healthy adult samples, and four of the NOD non-diabetic cluster with the diabetes samples (arrows).

### Ontogeny of β cell maturation and dedifferentiation

To distinguish, in an unbiased manner, between the reversal of ontogeny model and the Anna Karenina model of β cell dedifferentiation in diabetes, we generated a multi-dimensional trajectory map of the transcriptional states of β cells as they mature during development and as they lose their mature identity in each of the two types of diabetes (Figure 3). We reasoned that if the reversal of ontogeny model is correct, then diabetic β cells are expected to cluster along the developmental trajectory. On the other hand, if the Anna Karenina model is correct, then diabetic β cells will not cluster with any progenitor stage. Principal component analysis (PCA) of the top 15% most variable genes among the groups was used to generate a three-dimensional spatial distribution map of the samples. We found that the first three principal components captured 46.5% of the variation between the samples. PC1 (26.8% of the variation), PC2 (13.5% of the variation) and PC3 (6.2% of the variation) clearly separated the “healthy adult” samples, the “development” samples, and the “diabetic” samples into three distinct clusters (Figure 3, Left). Further separation was seen between the NOD-diabetic (T1D) and the BTBR-*Ob/Ob*-diabetic (T2D) samples (Figure 3, Right). Again, the NOD non-diabetic samples were divided between the NOD-diabetic and the healthy adult samples, indicating that loss of β cell maturation in NOD mice precedes the onset of overt diabetes. Thus, our analyses using two independent unsupervised mathematical methods suggest that β cells in the above two diabetes models lose their mature identity, but do not return to any developmentally relevant stage.

**Figure 3.**
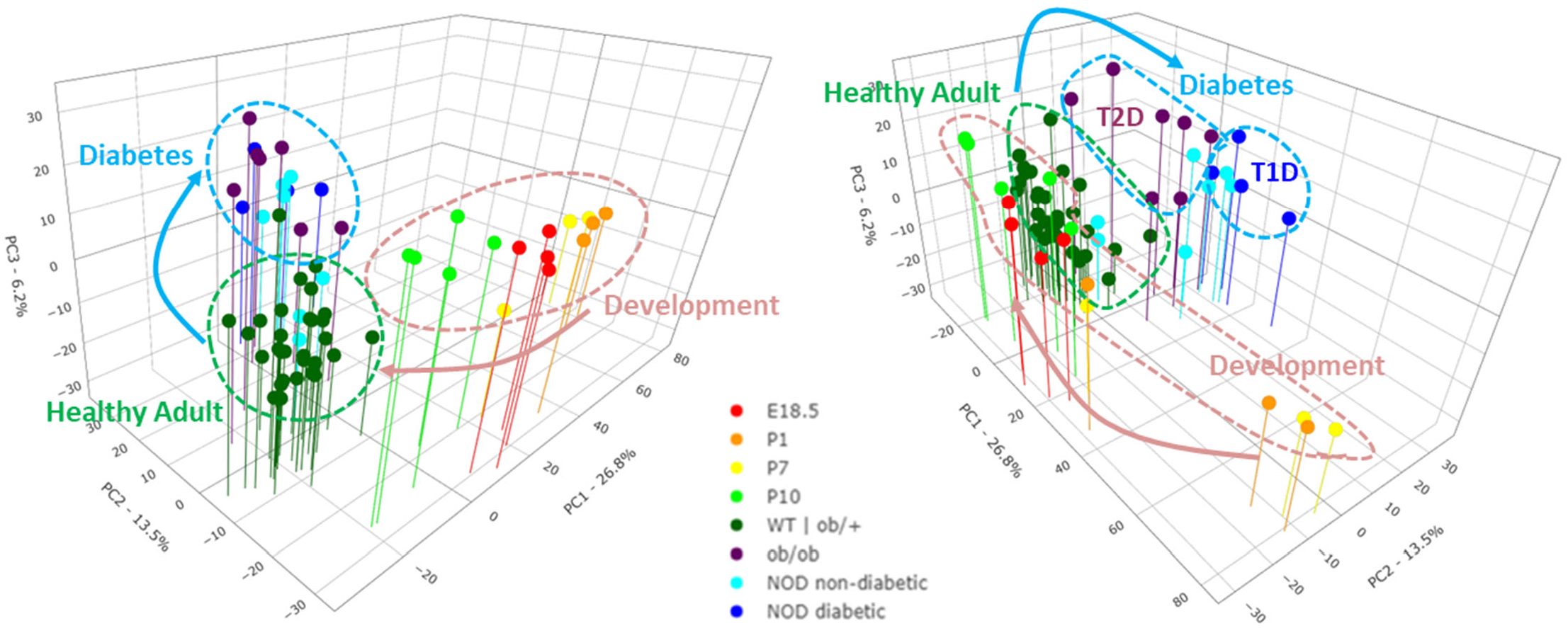
Different transcriptional trajectories in β cell maturation in development and their dedifferentiation in different types of diabetes. Principal component analysis of the top 15% most variable genes between β cell transcriptomes during development and dedifferentiation is shown. **Left:** β cells in diabetes cluster away from β cell of healthy adult mice, but do not cluster with any developmentally relevant stage, indicating that β cell dedifferentiation is not a reversal of developmental ontogeny. **Right:** View of the trajectory map from another angle, showing separation between β cell states in T1D and T2D.

### Gene-specific expression changes in β cell maturation and dedifferentiation

To validate our unbiased clustering results, we directly examined the expression of a broad list of published markers of mature β cell identity (20–29), “β cell disallowed” genes (30–32), markers of immature β cells and non-insulin-expressing β cell precursors (6, 8, 9, 20, 33–36), and islet hormones (Figure 4). Several markers of immature β cells and β cell progenitor genes (*MafB, Nnat, Sox17, Fev,* and *Myc*), as well as most “β cells disallowed” genes (*Ldha, Hk1, Mylk, Igfbp4, Ndrg2, Pcolce,* and *Slc16a2)* showed down-regulation during normal β cell maturation but were not re-expressed in either type of diabetes. One “disallowed” gene, *Ly6a*, was re-expressed in the T1D group and one disallowed gene, *Aldh1a3,* was re-expressed in the T2D group. Of the mature β cell genes, *MafA*, *Nkx6.1*, *Tshz1*, and *Slc2a2* were already present at high levels in the development group (which included semi-mature P10 pups (37)) and were down-regulated in the T2D group and, to a lesser extent, in the T1D group. Of the known β cell maturation markers, only *Ucn3* was up-regulated during β cell maturation and down-regulated in both types of diabetes, confirming previous reports by us and others that *Ucn3* is one of the most sensitive markers for the fully mature β cell state (18, 20, 38, 39). Conversely, *Dlk1,* a marker for immature β cells (20, 40), is down-regulated in maturation and is re-expressed in both types of diabetes, while *Gast* appears to be down-regulated in maturation and to be reexpressed specifically in T2D, as was previously reported (33). We did not see re-expression of *Neurog3* or any of the other markers of early β cell precursors in either type of diabetes.

**Figure 4.**
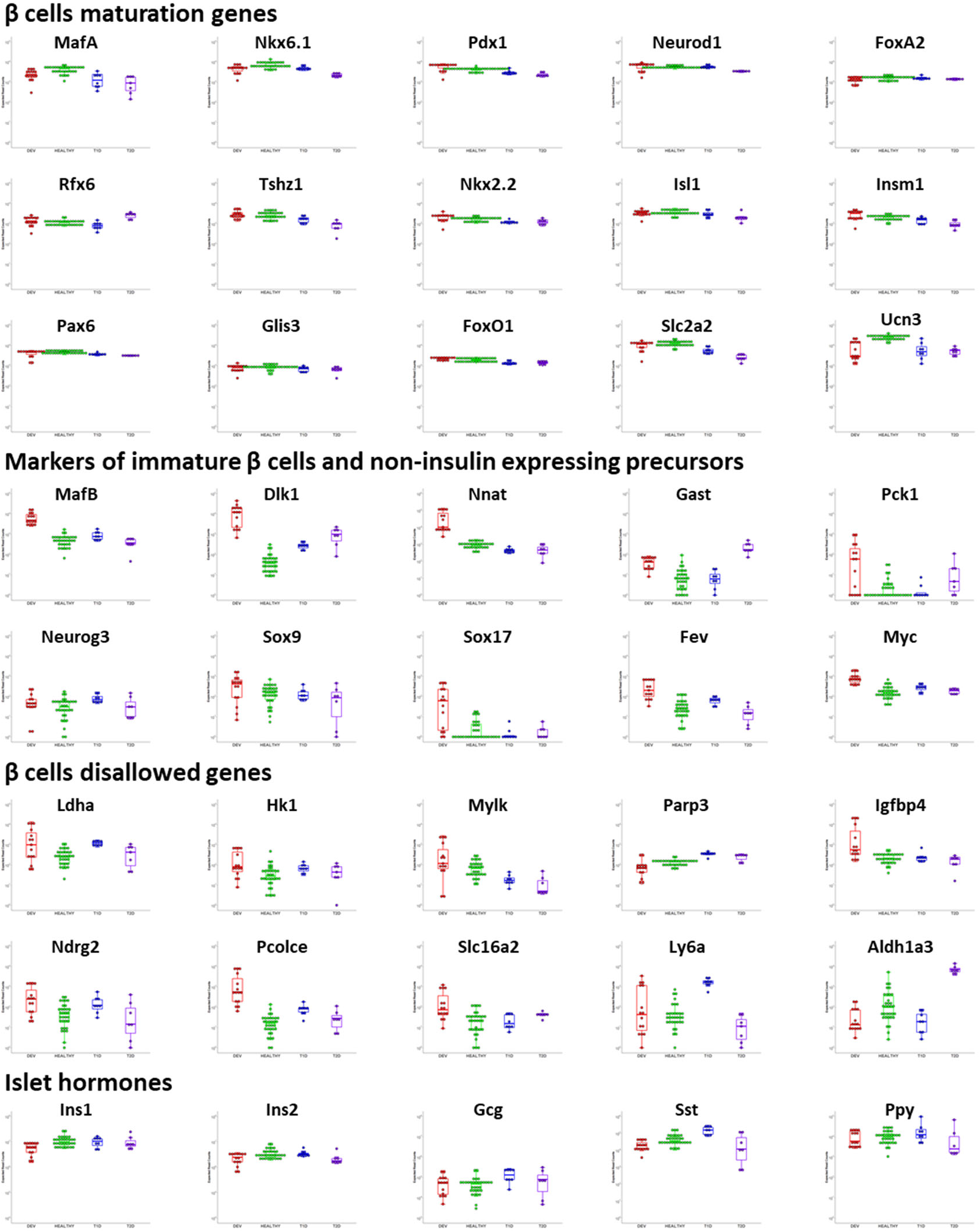
Expression of selected β cell maturation genes, β cell “disallowed” genes, markers of immature β cells and non-insulin expressing precursors, and islet hormones. Red: “development group”; green: “healthy adult” group; blue: “T1D-diabeic” group; purple: “T2D” group.

Further comparisons of all genes expressed at higher and lower than 1.5-fold with adjust p-value of less than 0.05 (*q*<0.05) in each non-mature condition (development, T1D-diabetic, and T2D-diabetic) compared to the healthy adult group showed little overlap among the non-mature groups, confirming our observation that dedifferentiated β cells in either of the diabetes groups do not revert to a developmentally relevant transcriptional state (Figure 5). A full list of genes in each group and Gene Ontology (GO) term enrichment of biological processes significantly enriched in each of the groups are presented in Supplementary Tables 1-4. Side by side comparisons of genes differentially expressed between each group and all other groups are shown in Supplementary Figure 1. These gene-specific analyses confirm distinct gene signatures for β cell maturation during normal postnatal development, and their dedifferentiation in each type of diabetes. We concluded that β cell dedifferentiation in diabetic NOD and BTBR-*Ob/Ob* mice is not a reversal of developmental ontogeny and is different for each type of diabetes.)

**Figure 5.**
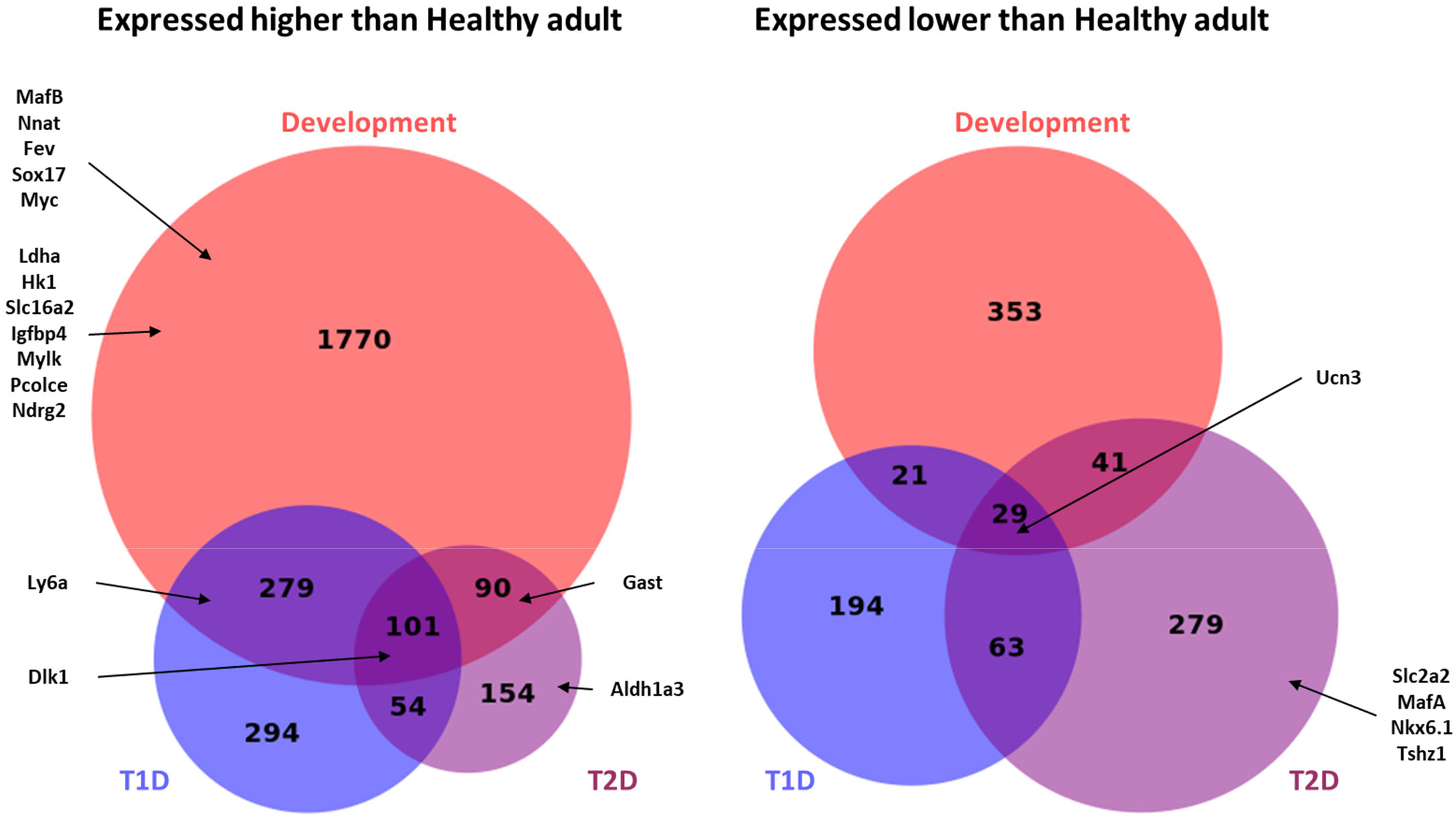
Comparison of gene-specific expression changes during β cell maturation and dedifferentiation in T1D and T2D. BioVenn diagram showing the number of genes up-regulated and down-regulated (1.5-fold, *q*<0.05) in each group compared to the healthy adult group. Representative genes enriched in each category are also presented. For a list of all genes in each group and enriched GO terms in each groups see Supplementary Tables 1-4.

## Discussion

Preventing or reversing β cell dedifferentiation is a promising approach to restoring glycemic control in people with diabetes. To this end, it is essential to understand the genetic mechanisms leading to β cell dedifferentiation under different diabetogenic conditions. Several studies over the last decade proposed that β cells in diabetes dedifferentiate in reverse order of their normal developmental ontogeny. This was shown by the loss of mature β cell markers in diabetic β cells, concomitant with the re-expression of several β cell progenitors genes, such as *Neurog3, Sox9, Myc,* and in some cases even *Nanog* and *Oct4* (8, 9, 34, 35). Other studies, however, reported the loss of mature β cell markers in diabetic β cells without re-expression of progenitor-stage transcription factors (16, 18, 41), or found that dedifferentiated β cells resemble immature (neonatal) β cells to some extent, but are not *Neurog3*-expressing progenitors (12, 33). It thus remains debated whether dedifferentiated β cells in diabetes revert to a progenitor-like state, or whether they lose their mature identity without reverting to any ontogeny-relevant stage and, if so, whether different types of diabetogenic stresses push β cells to different dedifferentiated trajectories. We set out to distinguish between the different models of β cell dedifferentiation in an unbiased manner, using unsupervised analysis of the transcriptional landscapes of both β cell maturation during development and their dedifferentiation in two mouse models of diabetes, namely NOD (a model for T1D) and BTBR-*Ob/Ob* (a model for T2D). We used the same lineage-tracing reporter system to isolate β cells both during development and during the progression to diabetes in the two different models of the disease. This allowed us to compare β cell maturation and their dedifferentiation in diabetes using one unperturbed system. We reasoned that superimposing the complete transcriptional states of the diabetic samples on the developmental ontogeny transcriptional map will directly resolve between the two models: if β cells in diabetes revert to any development-relevant transcriptional state, then the dedifferentiated samples will cluster along the developmental trajectory. On the other hand, if β cell dedifferentiation in diabetes is not a reversal of ontogeny, despite up-regulation of some genes that are expressed also in progenitors, then the dedifferentiated β cells will not cluster with any development-relevant stage. We report that, despite some similarities between immature and dedifferentiated β cells, such as reduced expression of several maturation markers and increased expression of some disallowed genes, β cells dedifferentiation, at least in the two mouse models tested here, is not a reversal of developmental ontogeny and is different between T1D and T2D.

It is worth noting that our analyses here focused on late (postnatal) β cell maturation. It is possible that if we compared our diabetic β cells to earlier (embryonic) progenitors, we would have found that there may be different entry points to β cell dedifferentiation, but the trajectories eventually converge to a stage resembling embryonic β cell precursors. However, we did not see re-expression of any known marker of early β cell percussors, including *Neurog3*, in either of the diabetes groups, even in samples from mice that were extremely diabetic for several weeks. Furthermore, our approach using an *Insulin* promoter-based genetic lineage-tracing system instead of a transient *Insulin* promoter-driven fluorescent reporter to isolate the cells means that we would have observed dedifferentiated β cells even if they were to drift into a non-insulin-expressing precursor state. This lineage-tracing system would have also detected β cell transdifferentiation into other endocrine and non-endocrine cell types, should that have been a substantial phenomenon. Another aspect of our approach that may confound our results is the possibility of contamination from small numbers of non-β cells, which are hard to sieve out using our bulk RNA-seq approach, such as mature acinar cells (few in early postnatal pancreata and increasing in adults), immune cells (higher probability of contamination in T1D samples), or adipose cells (more abundant in samples from obese-diabetic mice). Indeed, several genes associated with such contamination are detected in our comparisons. While such contamination may possibly skew our unsupervised clustering analyses to some extent, our FACS-sorting using lineage-traced β cells and our bulk RNA-seq approaches compensate for such rare events due to the analyses being done on relatively purified β cell populations, and the depth of sequencing, which is not possible with single-cell RNA-seq. Most importantly, our gene-specific analyses using a large list of known markers of β cell development and maturation provide independent confirmation that dedifferentiated β cells in diabetic NOD and BTBR-*Ob/Ob* mice lose their mature β cell identity, but do not return to any developmentally-relevant state. That said, our results do not dispute that β cells in other models not tested here, such as *FoxO!*-null mice (8) and mice subjected to a fasting-mimicking diet (42) could return to a *Neurog3*-expressing progenitor state. With Neurog3 being a master regulator of an embryonic proto-endocrine transcriptional program (43, 44), it is conceivable its re-expression in these unique models may force a more developmentally-relevant cell identity that is not seen in β cells from diabetic NOD or BTBR-*Ob/Ob* mice.

We propose that at least in the case of diabetic NOD and BTBR-*Ob/Ob* mice, each type of diabetes produces β cells that are dedifferentiated in their own way, supporting the Anna Karenina model of β cell dedifferentiation. We hope that these results will provide a valuable resource in the efforts of finding genetic and pharmacological intervention points for preventing and possibly reversing β cell dedifferentiation in diabetes.

## Methods

### Mice

All animal experiments were conducted in accordance with the University of Wisconsin-Madison IACUC guidelines under protocol number M005221. *BTBR-Lep^ob/ob^,* NOD, and ICR (*“WT”)* mice were obtained from the Jackson Laboratories and Envigo. *Insulin2-Cre;Rosa26-lox-stop-lox-H2BmCherry* mice were previously reported (18). Blood glucose and weight were measured in non-fasted animals using OneTouch Ultra2 glucometer (LifeScan, Milpitas, CA) at the animal facility before islet collection. Mice with blood glucose higher than 300 mg/dL were considered as diabetic. Islet isolation was performed as previously described (18, 20). Isolated islets were dissociated with 0.25% trypsin-EDTA before sorting through BD FACS Aria II for mCherry+ cells.

### RNA sequencing

RNA was isolated from FACS sorted lineage-traced β cells from ICR embryos, neonates, and adult mice; NOD adult (diabetic and non-diabetic) mice, and BTBR-*Ob/Ob* and BTBR-*Ob/+* adult mice using phenol chloroform extraction (TRIzol) and Qiagen RNeasy Plus Mini Kit (Qiagen). DNA libraries were generated using Takara’s SMART-Seq v4 Low Input RNA Kit for Sequencing (Takara, Mountain View, California, USA) for cDNA synthesis and the Illumina NexteraXT DNA Library Preparation (Illumina, San Diego, CA, USA) kit for cDNA dual indexing. Full length cDNA fragments were generated from 1-10ng total RNA by SMART (Switching Mechanism at 5’ End of RNA Template) technology and were sequenced for 1×100 on Illumina NovaSeq 6000 at sequencing depth of 25-30 M reads per sample. Two samples (1 BTBR-*Ob/Ob* and 1 BTBR-*Ob/+*) were sequenced for 2×100 on Illumina NovaSeq 6000 at the same sequencing depth. Quality control (QC) of both single-end and paired-end raw sequencing data was conducted using FastQC-0.11.7 (45) and MultiQC-1.9 (46). All samples passed QC as they have uniformly high base quality and sequence quality. Mild adapter contaminations were detected, and we decided not to perform adapter trimming. Raw sequencing data were aligned to mm10 reference genome using Bowtie-1.2.2 (47) under default settings. Gene-by-sample count matrix was estimated using RSEM-1.3.0 (48) under default settings. After combining the two batches into one dataset, genes with average expression less than 1 were filtered out. Median-by-ratio normalization (49) was conducted on combined data to account for sequencing depth artifact and batch effects. This results in a normalized expression matrix with 63 samples and 16455 genes. All sequencing data are available in the Gene Expression Omnibus (GEO) repository under accession number ########.

### Hierarchical clustering and principal component analysis

The normalized expression matrix was log2 transformed and further adjusted for potential batch effects by removeBatchEffect() in the limma R package (50) (v3.44.3). The 15% most variable genes were identified using varFilter() in the genefilter R package (v1.70). Hierarchical clustering was performed on these highly variable genes using Spearman correlational distance and Ward’s linkage method in the cluster R package (51) (v2.1) and visualized on dendextend (52) R packages (v1.14). For principal component analysis, eigenvectors were calculated using the prcomp() function, and 3D visualization was generated by the Plotly R package (v4.9.2.1).

### Differential Expression and fold change

Genes having non-zero expression in 10 or more samples and at least 5 reads total were retained for differential expression (DE) analysis. DESeq2 (53) (v1.28.1) was used to identify DE genes. Specifically, we applied DESeq2 to obtain gene-specific *p*-values which were converted to *q*-values using the Benjamini and Hochberg method. A gene was considered DE if its *q*-value <0.05 and if its shrunken log2 fold change, estimated using lfcShrink() in the DESeq2 package, exceeded 1.5. Visualization was done in Biovenn (54) and the EnchancedVolcano R package (v1.6).

## Author contributions

Conceptualization, B.B. and S.D.N.; Methodology, B.B., S.D.N., and C.K.; Data Acquisition, S.D.N.; Formal Analysis, S.D.N, Z.N., and J.B.; Writing Original Draft, B.B. and S.D.N.; Writing, Review and Editing, all authors; Funding Acquisition, B.B.; Supervision, B.B. and C.K.

## Acknowledgments

We thank N. Sharon, D. Ben-Zvi, A. Helman, and A. Attie for valuable comments and discussion. We are grateful to all present and former members of the Blum Lab, especially Melissa Adams, Jennifer Gilbert, Bayley Waters, Emily Cade, Ron Fleminger, and Emily Maritato for help on this project. We are also grateful to the University of Wisconsin-Madison Biotechnology Center Gene Expression Core for RNA sequencing. This work was funded in part by grants number 1R56DK115837 from the NIDDK and 2-SRA-2018-621-S-B from the JDRF to BB, and grant number 1S10RR025483-01 to the University of Wisconsin-Madison School of Medicine and Public Health Flow Cytometry Core.

---

**Supplementary Figure 1.**
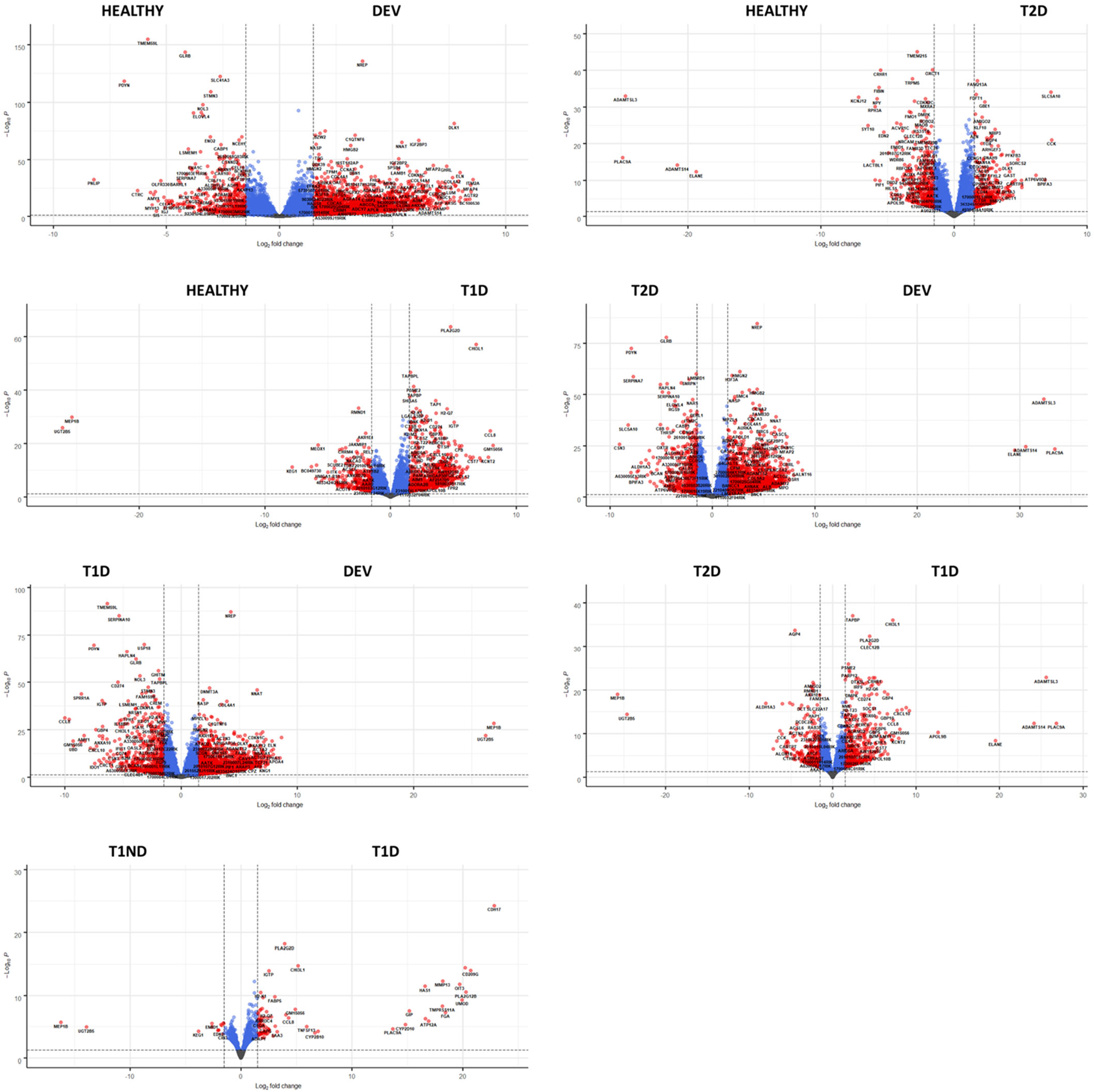
Gene-specific expression changes between each of the groups and all other groups. Shown are volcano plots of genes differentially expressed between each group compared to all other groups.

